# Every single conidium in *Aspergillus fumigatus* caspofungin tolerant strains are intrinsically caspofungin tolerant

**DOI:** 10.1101/2022.02.17.480978

**Authors:** Clara Valero, Ana Cristina Colabardini, Patrícia Alves de Castro, Jorge Amich, Michael J. Bromley, Gustavo H. Goldman

**Author notes:** Corresponding author: Dr. Gustavo Henrique Goldman, Faculdade de Ciências Farmacêuticas de Ribeirão Preto, Universidade de São Paulo, Av. do Café S/N CEP 14040-903, Ribeirão Preto, São Paulo, Brazil.

## Abstract

*Aspergillus fumigatus* is a human fungal pathogen that causes a disease named aspergillosis. Echinocandins, such as the fungistatic drug caspofungin (CAS) are used as second-line therapy. Some *A. fumigatus* clinical isolates can survive and grow in higher CAS concentrations, a phenomenon known as “caspofungin paradoxical effect” (CPE). Here we investigate if CPE is due to a subpopulation of conidia produced by a CAS tolerant strain, indicative of a persistence phenotype or is caused by all the conidia which would be consistent with a tolerance phenotype. We evaluated 67 *A. fumigatus* clinical isolates for CPE growth and used a novel CPE Index (CPEI) classified them as CPE^+^ (CPEI ≥ 0.40) or CPE^−^ (CPEI ≤ 0.20). Conidia produced by three CPE^+^ clinical isolates, CEA17 (CPEI=0.52), Af293 (CPEI=0.64), CM7555 (CPEI=0.58) all showed the ability to grow in high levels of CAS while all conidia produced by the CPE^−^ isolate IFM61407 (CPEI=0.12) strain showed no evidence of tolerance. Given the importance of calcium/calcineurin/transcription factor CrzA pathway in CPE regulation, we also evaluated Δ*crzA*^Af293^ (CPE^−^) and Δ*crzA*^CEA17^ (CPE^+^) conidia tolerance to CAS. All Δ*crzA*^CEA17^ conidia showed CPE^+^ while 100 % of Δ*crzA*^Af293^ spores are CPE^−^. As all spores derived from an individual strain are phenotypically indistinct with respect to CPE it is likely that CPE is a genetically encoded adaptive trait that should be considered an antifungal tolerant phenotype. As the CPEI shows that the strength of the CPE is not uniform between strains we propose that the mechanisms that govern this phenomenon are multi-factorial.

## Importance

*Aspergillus fumigatus* is the most important human fungal pathogen causing pulmonary infections. Caspofungin (CAS) is a fungistatic drug used as a second-line therapy against aspergillosis, the group of diseases caused by *A. fumigatus*. CAS inhibits the function of Fks1 a β-1,3-glucan synthase that has a critical role in the synthesis of the cell wall. Resistance to CAS is commonly associated with mutations in *fks1*, however, some *A. fumigatus* clinical isolates are able to grow in the presence of higher CAS concentrations, a drug tolerance phenomenon known as the “caspofungin paradoxical effect” (CPE). Here, based on the characterization of CPE presence in a series of *A. fumigatus* clinical isolates, we demonstrate that *A. fumigatus* CAS tolerant strains do not exhibit a CAS heterogeneous response. Our results indicate that every single conidium in *A. fumigatus* CAS tolerant strains are intrinsically CAS tolerant.

## Observation

Fungal diseases are a significant health problem affecting more than 1 billion people around the world and causing more than 1.5 million deaths (1). *Aspergillus fumigatus* is the most important agent of fungal pulmonary infection causing a group of heterogeneous clinical diseases named aspergillosis (2). Few antifungal agents, such as the fungicidal azoles (first-line therapy, itraconazole, posaconazole, voriconazole, and isavuconazole), amphotericin B, and the fungistatic echinocandins (caspofungin, CAS, second-line therapy) are available to treat aspergillosis while worryingly clinical azole resistance has been increasingly reported (3–5). While azoles inhibit the ergosterol biosynthesis pathway by directly targeting the eburicol 14-demethylase Cyp51A/ERG11 (6), CAS acts by noncompetitively inhibiting the fungal β-1,3-glucan synthase (Fks1), which is essential for the biosynthesis of β-1,3-glucan in the fungal cell wall (7). In patients suffering from invasive aspergillosis, strains resistant to the azoles have often been shown to be acquired from the environment however in those suffering from chronic forms of aspergillosis resistance typically, and frequently occurs during the course of infection (8). CAS resistance has been increasingly observed in *Candida* spp. (9), and although infrequently described, there are reports of *A. fumigatus* CAS resistance from patients with chronic aspergillosis (10). Mutations in specific “hotspots” of the FKS1 gene are the main genetic mechanisms of CAS resistance described in both *Candida* spp. and *Aspergillus* spp. (10, 11).

Drug tolerance has been extensively studied in bacterial pathogens, where it has been defined as the ability of all cells of an isogenic strain to survive and even grow at slow rates in the presence of drug concentrations that are greater than the minimal inhibitory concentration (MIC) (12). The term persistance is a type of tolerance that describes a phenomenon where only a sub-population of cells within an isogenic strain are tolerant. To date, the description of tolerance in fungi has focused almost exclusively on yeast-like fungi where tolerance is frequently observed as occurring in sub-populations within an isogenic strain (13–15). Very little attention has been given to defining drug tolerance in filamentous fungi however one adaptive phenomenon exhibited by some *A. fumigatus* clinical isolates allows them to grow and tolerate high CAS concentrations and is known as the “caspofungin paradoxical effect” (CPE). Although there are several mechanisms already described for *A. fumigatus* CPE (for reviews, see 16, 17), there is a little understanding if CPE occurs as a result of phenotypic heterogeneity within an isogenic population. Here, based on the characterization of CPE presence in a series of *A. fumigatus* clinical isolates, we demonstrate that conidia from *A. fumigatus* CAS tolerant strains do not exhibit CAS heterogeneity and hence CPE should be considered as a tolerant but not persistant phenotype.

We have investigated the CPE in 67 *A. fumigatus* clinical isolates (**Supplementary Tables S1 and S2 at 10.6084/m9.figshare.19178888;** S. Zhao, A. Martin-Vicente, A. C. Colabardini, L. P. Silva, J. R. Fortwendel, G. H. Goldman, and J. G. Gibbons, submitted for publication) and defined a CPE index (CPEI) = average radial diameter in MM+8 µg/ml CAS/average radial diameter in MM, where CPEI ≥ 0.4 are CPE^+^, CPEI ≥ 0.25 and ≤ 0.4 have an intermediate (partial) phenotype (CPE^P^), and CPEI ≤ 0.2 are CPE^−^. **Figure 1A** shows the heat map representing the CPEI of 67 *A. fumigatus* clinical isolates grown for 5 days at 37 °C on MM+0.125 to 8 µg/ml of CAS. Radial growth in the presence of CAS is exemplified for three clinical isolates CEA17/A1163 (CPEI=0.52), CM7555 (CPEI=0.58) and IFM61407 (CPEI=0.12) (**Figure 1B**). It should be noted that the CPEI cannot be assessed for strains that are resistant to CAS using the drug concentrations outlined here however by adjusting the CAS concentrations the CPEI can still be calculated.

**Figure 1.**
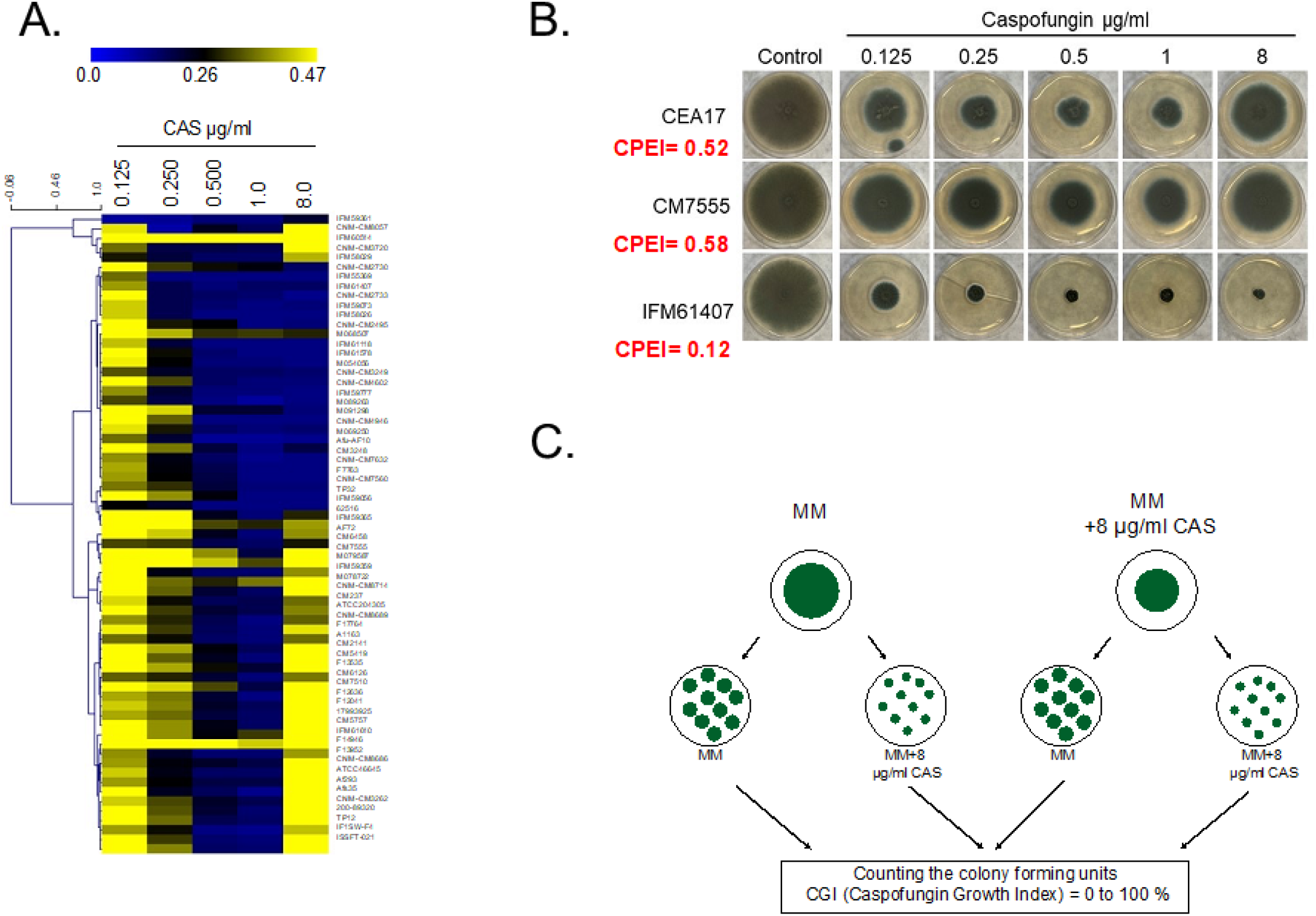
Distribution of *A. fumigatus* CAS tolerance in 67 clinical isolates and definition of CPE index (CPEI) and CAS Growth Index (CGI). **(A)** Heat map depicting the CPE Index (CPEI) = radial diameter in MM+8 µg/ml CAS/radial diameter in MM, where CPEI ≥ 0.4 are CPE^+^, CPEI ≥ 0.25 and ≤ 0.4 are partially CPE^+^ (CPE^P^), and CPEI ≤ 0.2 are CPE^−^. Heat map scale and gene identities are indicated. Hierarchical clustering was performed in MeV (http://mev.tm4.org/), using Pearson correlation with complete linkage clustering. **(B)** Growth of *A. fumigatus* CEA17, CM7555, and IFM61407 clinical isolates on MM and MM+CAS (increasing concentrations). The strains were grown for 5 days at 37 °C. **(C)** Scheme showing how the CAS Growth Index (CGI) was calculated. *A. fumigatus* isolates were grown on MM or MM+8 µg/ml for 5 days at 37 °C. Conidia were harvested, filtered, diluted to 10^3^ sp/ml and 100 µl were plated in MM or MM+8 µg/ml and incubated for 2 or 3 days at 37 °C. The number of colonies were counted in both treatments and determined the CGI as follows: CGI (%) = (number of colonies radial diameter ≥0.5 cm on MM+8 µg/ml CAS/number of colonies radial diameter ≥0.5 cm MM) x 100.

*A. fumigatus* sexual and asexual spores are the single developmental “cell-like”-structures with a single nucleus in the fungus. Germlings or mycelia are syncytia with several nuclei present in a common cytoplasm. Is CAS tolerance present in a “fraction” of the conidial population or in every single conidium in a single CAS-tolerant CPE^+^ clinical isolate? In order to address this question, we have grown two *A. fumigatus* reference isolates CEA17 (CPEI=0.52) and Af293 (CPE=0.64) in MM in the presence or absence of CAS 8µg/ml for 5 days at 37 °C. Then, conidia were harvested, filtered, diluted to 10^3^ conidia/ml and 100 µl were plated them on MM (control) or MM+8 µg/ml (a CPE concentration) (**Figure 1D**). After 48 hs (MM) or 72 hs (MM+CAS) of growth at 37 °C, the number of colonies ≥ 0.5 cm radial diameter were counted (**Figure 1C**), and the CAS growth index, CGI (%) = (number of colonies radial diameter ≥0.5 cm on MM+8 µg/ml CAS/number of colonies radial diameter ≥0.5 cm MM) x 100, was determined (**Figure 1C**). When CEA17, Af293, and CM7555 clinical isolates were grown on either MM or MM+8.0 µg/ml, we observed a CGI of 100 % (**Figure 2A, Supplementary Table S3 at 10.6084/m9.figshare.19178888**). We did not observe radial diameter size heterogeneity in any of the colonies grown on MM or MM+8 µg/ml (all c. 0.5 cm radial diameter; **Figure 2A**). These results indicate that every single conidium in *A. fumigatus* CPE^+^ strains are intrinsically CAS tolerant. We then evaluated the CGI for the clinical isolate IFM61407 strain that is CPE^−^ (**Figures 1C**). IFM61407 conidia derived from MM or MM+8.0 µg/ml CAS show a CGI of 0 %, respectively (**Figure 2A, Supplementary Table S3 at 10.6084/m9.figshare.19178888**).

**Figure 2.**
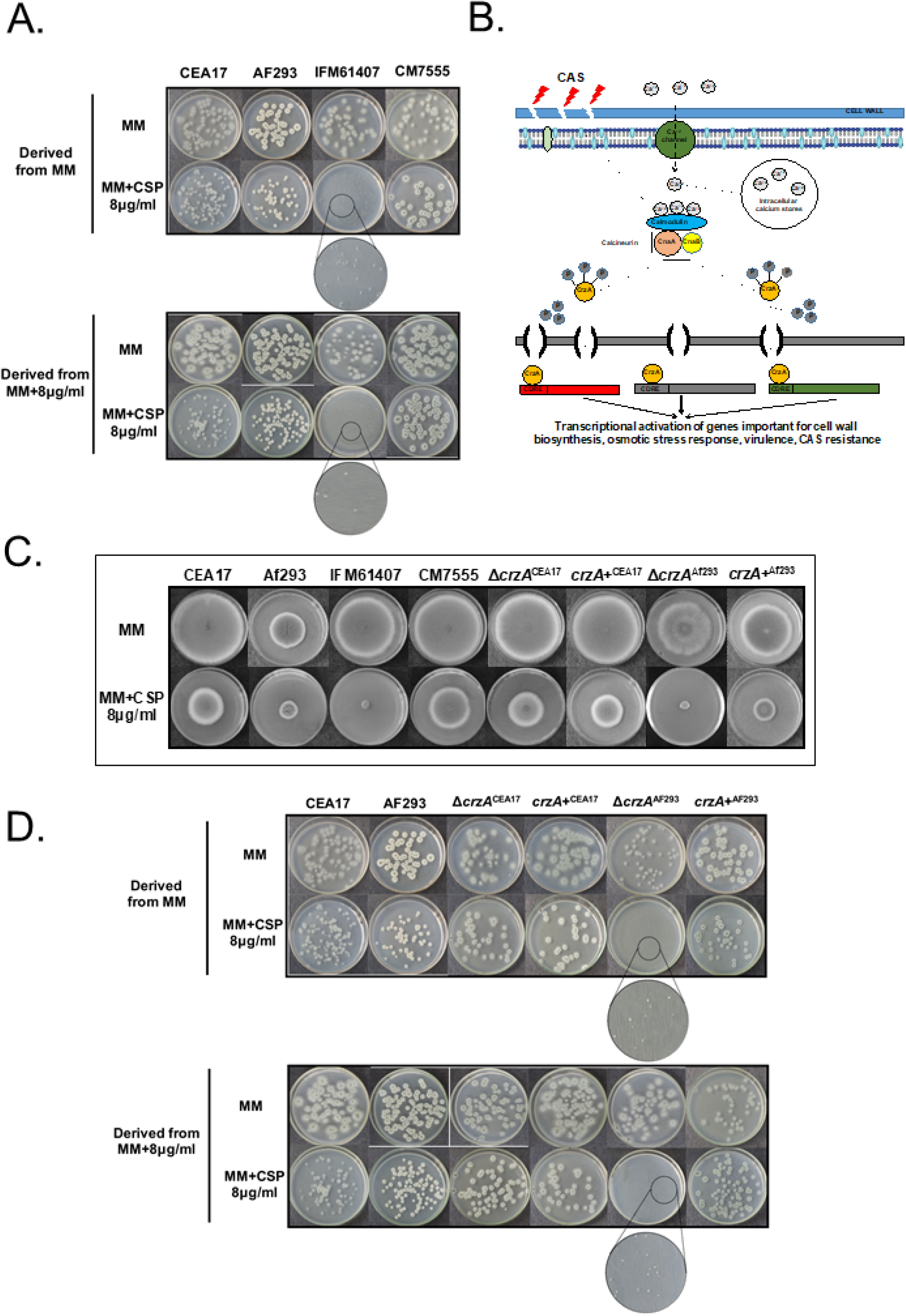
CAS growth index (CGI) for *A. fumigatus* clinical isolates. **(A)** CEA17, Af293, CM7555, and IFM61407 clinical isolates were grown on MM and MM+8 µg/ml of CAS for 5 days at 37 °C. Conidia were harvested, filtered, diluted to 10^3^ sp/ml and 100 µl were plated in MM or MM+8 µg/ml and incubated for 2 or 3 days at 37 °C. The number of colonies were counted in both treatments and the CGI was determined. **(B)** Scheme showing the calcium/calcineurin/CrzA pathway. Upon CAS cell wall damage, calcium concentrations increase in the cytoplasm by calcium transport or mobilization of endogenous calcium deposits. Calcium binds to calmodulin, which activates calcineurin that directly dephosphorylates CrzA, resulting in its translocation to the nucleus. CrzA binds to calcineurin-dependent response elements (CDRE) promoters, activating the transcriptional programs that promote stress tolerance. **(C)** CEA17, Af293, Δ*crzA*^CEA17^, and Δ*crzA*^Af293^ strains were grown on MM and MM+CAS for 5 days at 37 °C. **(D)** Δ*crzA*^CEA17^ and Δ*crzA*^Af293^ strains were grown on MM and MM+CAS for 5 days at 37 °C. Conidia were harvested, filtered, diluted to 10^3^ sp/ml and 100 µl were plated in MM or MM+8 µg/ml and incubated for 2 or 3 days at 37 °C. The number of colonies were counted in both treatments and the CGI was determined.

Calcium homeostasis has been reported to play a central role in the CPE cellular response in *A. fumigatus* (18, 19, **Figura 2B**). CAS increases the intracellular calcium (Ca^2+^) concentration activating the calcineurin-CrzA pathway (20). CrzA regulates the activation of several stress responses and cell wall modifications (21, 19). Interestingly, *crzA* deletion in the clinical strain Af293 results in CPE loss (22, **Figure 2C**), while *crzA* deletion in the CEA17 background is CPE^+^ (23, 19, **Figure 2C**), demonstrating intra-species differences or CPE heterogeneity. Differently from Δ*crzA*^CEA17^ mutant, the Δ*crzA*^Af293^ mutant cannot activate cell wall remodeling genes upon CAS exposure, affecting its CPE (23). The CGIs for Δ*crzA*^CEA17^ and Δ*crzA*^Af293^ mutant strains are 100 and 0 %, respectively, when grown on MM+8 µg/ml independently if the conidia are derived from MM or MM+8 µg/ml (**Figure 2D, Supplementary Table S3 at 10.6084/m9.figshare.19178888**). Taken together these results indicate that the transcription factor CrzA, whose deletion results in heterogeneity in the response of the CEA17 and Af293 strains to CAS does not show CPE heterogeneity since all the conidia from the CPE^−^ Δ*crzA*^Af293^ strain do not exhibit CPE, while all the conidia from the CPE^+^ Δ*crzA*^CEA17^ strain are CPE^+^ (**Figure 2D**).

Our results emphasize the view that every single conidium in an *A. fumigatus* CPE strain is able to grow and be CAS tolerant. In contrast, conidia from *A. fumigatus* strains lacking CPE have reduced growth in CPE concentrations, strongly indicating that there are no *A. fumigatus* CAS tolerant subpopulations. As a conclusion, *A. fumigatus* CPE is a homogeneous trait within an isogenic population while CPE heterogeneity exists between strains.

## Acknowledgements

We thank Fundação de Amparo à Pesquisa do Estado de São Paulo (FAPESP) 2018/00715-3 (CV), 2017/07536-4 (ACC), 2016/12948-7 (PAC) and 2016/07870-9 (GHG), and Conselho Nacional de Desenvolvimento Científico e Tecnológico (CNPq) 301058/2019-9 and 404735/2018-5 (GHG), both from Brazil, and National Institutes of Health/National Institute of Allergy and Infectious Diseases (R01AI153356), from the USA. This work was also supported by the Wellcome Trust grant number 219551/Z/19/Z and 208396/Z/17/Z to M.B.

## References

1. Brown GD, Denning DW, Gow NA, Levitz SM, Netea MG, White TC. 2012. Hidden killers: human fungal infections. Sci Transl Med 4:165rv13.

2. Arastehfar A, Carvalho A, Houbraken J, Lombardi L, Garcia-Rubio R, Jenks JD, Rivero-Menendez O, Aljohani R, Jacobsen ID, Berman J, Osherov N, Hedayati MT, Ilkit M, Armstrong-James D, Gabaldon T, Meletiadis J, Kostrzewa M, Pan W, Lass-Florl C, Perlin DS, Hoenigl M. 2021. Aspergillus fumigatus and aspergillosis: From basics to clinics. Stud Mycol 100:100115.

3. Ostrosky-Zeichner L, Al-Obaidi M. 2017. Invasive fungal infections in the intensive care unit. Infect Dis Clin North Am 31:475–487.

4. Perfect JR. 2017. The antifungal pipeline: a reality check. Nat Rev Drug Discov 16:603–616.

5. Robbins N, Caplan T, Cowen LE. 2017. Molecular evolution of antifungal drug resistance. Annu Rev Microbiol 71:753–775.

6. Warrilow AG, Parker JE, Price CL, Nes WD, Kelly SL, Kelly DE. 2015. In vitro biochemical study of CYP51-mediated azole resistance in Aspergillus fumigatus. Antimicrob Agents Chemother 59:7771–8.

7. Perlin DS. 2015. Mechanisms of echinocandin antifungal drug resistance. Ann N Y Acad Sci 1354:1–11.

8. Garcia-Rubio R, Cuenca-Estrella M, Mellado E. 2017. Triazole resistance in Aspergillus species: an emerging problem. Drugs 77:599–613.

9. Shields RK, Nguyen MH, Clancy CJ. 2015. Clinical perspectives on echinocandin resistance among Candida species. Curr Opin Infect Dis 28:514–22.

10. Jimenez-Ortigosa C, Moore C, Denning DW, Perlin DS. 2017. Emergence of echinocandin resistance due to a point mutation in the fks1 gene of Aspergillus fumigatus in a patient with chronic pulmonary aspergillosis. Antimicrob Agents Chemother 61 (12):e01277–17.

11. Wiederhold NP. 2016. Echinocandin resistance in Candida species: a review of recent developments. Curr Infect Dis Rep 18:42.

12. Brauner A, Fridman O, Gefen O, Balaban NQ. 2016. Distinguishing between resistance, tolerance and persistence to antibiotic treatment. Nat Rev Microbiol 14:320–30.

13. Delarze E, Sanglard D. 2015. Defining the frontiers between antifungal resistance, tolerance and the concept of persistence. Drug Resist Updat 23:12–19.

14. Berman J, Krysan DJ. 2020. Drug resistance and tolerance in fungi. Nat Rev Microbiol 18:319–331.

15. Ben-Ami R, Kontoyiannis DP. 2021. Resistance to antifungal drugs. Infect Dis Clin North Am 35:279–311.

16. Aruanno M, Glampedakis E, Lamoth F. 2019. Echinocandins for the treatment of invasive aspergillosis: from laboratory to bedside. Antimicrobial agents and chemotherapy 63(8) :e00399–19.

17. Papon N, Morio F, Sanglard D. 2020. Signaling pathways governing the caspofungin paradoxical effect in Aspergillus fumigatus. mBio 11(4) :e01816–20.

18. Juvvadi PR, Munoz A, Lamoth F, Soderblom EJ, Moseley MA, Read ND, Steinbach WJ. 2015. Calcium-mediated induction of paradoxical growth following caspofungin treatment is associated with calcineurin activation and phosphorylation in Aspergillus fumigatus. Antimicrobial agents and chemotherapy 59(8):4946–55.

19. Ries LNA, Rocha MC, de Castro PA, Silva-Rocha R, Silva RN, Freitas FZ, de Assis LJ, Bertolini MC, Malavazi I, Goldman GH. 2017. The Aspergillus fumigatus CrzA transcription factor activates chitin synthase gene expression during the caspofungin paradoxical effect. mBio 8 (3):e00705–17.

20. Park HS, Lee SC, Cardenas ME, Heitman J. 2019. Calcium-calmodulin-calcineurin signaling: a globally conserved virulence cascade in eukaryotic microbial pathogens. Cell Host Microbe 26:453–462.

21. Soriani FM, Malavazi I, da Silva Ferreira ME, Savoldi M, Von Zeska Kress MR, de Souza Goldman MH, Loss O, Bignell E, Goldman GH. 2008. Functional characterization of the Aspergillus fumigatus CRZ1 homologue, CrzA. Mol Microbiol 67:1274–91.

22. Fortwendel JR, Juvvadi PR, Perfect BZ, Rogg LE, Perfect JR, Steinbach WJ. 2010. Transcriptional regulation of chitin synthases by calcineurin controls paradoxical growth of Aspergillus fumigatus in response to caspofungin. Antimicrobial agents and chemotherapy. 54(4):1555–63.

23. Colabardini AC, Wang F, Dong Z, Pardeshi L, Rocha MC, Costa JH, Dos Reis TF, Brown A, Jaber QZ, Fridman M, Fill T, Rokas A, Malavazi I, Wong KH, Goldman GH. 2022. Heterogeneity in the transcriptional response of the human pathogen Aspergillus fumigatus to the antifungal agent caspofungin. Genetics 220 (1):iyab183.

